# Beta-3 Adrenoreceptors protect from hypertrophic remodelling through AMP-Activated Protein Kinase and Autophagy

**DOI:** 10.1101/2020.01.14.905893

**Authors:** Emilie Dubois-Deruy, Roselle Gelinas, Christophe Beauloye, Hrag Esfahani, Chantal Dessy, Luc Bertrand, Jean-Luc Balligand

## Abstract

**Aims:** The abundance of beta3-adrenergic receptors (β3-ARs) is upregulated in diseased human myocardium. We previously showed that cardiac-specific expression of β3-AR inhibits the hypertrophic response to neurohormonal stimulation. Here, we further analyzed signalling pathways involved in the anti-hypertrophic effect of β3-AR.

**Methods:** *In vitro* hypertrophic responses to phenylephrine (PE) were analyzed in neonatal rat ventricular myocytes (NRVM) infected with a recombinant adenovirus expressing the human β3-AR (AdVhβ3). We confirmed results in mice with cardiomyocyte-specific moderate expression of human β3-AR (β3-TG) and WT littermates submitted to thoracic transverse aortic constriction (TAC) for 9 weeks.

**Results:** We observed a colocalization of β3-AR with the AMP-activated protein kinase (AMPK) both in neonatal rat andin adult mouse cardiomyocytes. Treatment of NRVM with PE induced hypertrophy and a decrease in phosphorylation of Thr172-AMPK (/2, p=0.0487) and phosphorylation of Ser79- acetyl-CoA carboxylase (ACC) (/2.6, p=0.0317), inducing an increase in phosphorylated Ser235/236 S6 protein (x2.5, p=0.0367) known to be involved in protein synthesis. These effects were reproduced by TAC in WT mice, but restored to basal levels in β3-AR expressing cells/mice. siRNA targeting of AMPK partly abrogated the anti-hypertrophic effect of β3-AR in response to PE in NRVM (x1.3, p<0.0001). Concomitant with hypertrophy, autophagy measured by microtubule-associated protein 1 light chain 3 (LC3)-II/LC3-I ratio and p62 abundance was decreased by PE in NRVM (/2.6, p=0.0010 and x3, p=0.0016, respectively) or TAC in WT mice (/5.4, p=0.0159); and preserved in human β3-AR expressing cells and mice, together with reduced hypertrophy.

**Conclusions:** Cardiac-specific moderate expression of β3-AR inhibits the hypertrophic response in part through AMPK activation followed byinhibition of protein synthesis and preservation of autophagy. Activation of the cardiac β3-AR pathway may provide future therapeutic avenues for the modulation of hypertrophic remodelling.

## INTRODUCTION

Haemodynamic overload, ischemic or oxidative stress promote adverse cardiac remodelling, a leading cause of worsening heart failure^1^. Most of these pathophysiologic conditions are associated with adrenergic stimulation, resulting in adrenoceptor (AR) activation on different cell types within the myocardium. Among these, β1-AR’s are classically considered to mediate short-term positive effects on all aspects of myocardial contractility; however, long-term stimulation produces adverse effects on myocardial remodelling, in part through activation of calcium-dependent pro-hypertrophic effects, ultimately associated with cardiomyocyte loss^2^. Such “maladaptive” remodelling is usually accompanied with left ventricular (LV) geometry disruption and interstitial fibrosis leading to progressive diastolic and systolic heart failure. Conversely, β3-AR mediates effects on cardiac myocytes that are antipathetic to β1-AR, i.e. protection from hypertrophic^3^ or fibrotic^4^ remodelling. Deciphering the underlying signalling pathways should yield new therapeutic strategies to favourably modulate remodelling.

AMP-activated protein kinase (AMPK) is a heterotrimeric serine/threonine protein kinase composed of one catalytic (AMPKα) and two regulatory subunits (AMPKβ and AMPKγ) present under several isoforms (two for α and β and three for γ)^5^. AMPK is an important cellular sensor of energy level. Its activation derives from an increase in the intracellular AMP-to-ATP ratio that typically occurs during metabolic stresses, such as myocardial ischemia^6^. AMP stimulates AMPK allosterically via its binding to the AMPK γ-subunit and protects AMPK from de-phosphorylation on Thr172, a residue located in the activation loop of the AMPKα catalytic subunit. Once activated, AMPK participates in the maintenance of energy homeostasis by switching off anabolic pathways that consume ATP and switching on alternative catabolic pathways that generate ATP^7^. It is currently recognized that AMPK effects extend beyond its metabolic role^7^. In agreement, it has been shown that AMPK activation prevents cardiac hypertrophy development^7^. Accordingly, AMPK deletion leads to enhanced hypertrophy^8^. At the molecular level, AMPK is known to inhibit the mechanistic target of rapamycin (mTOR) pathway via a complex mechanism^7^. The mTOR pathway promotes protein synthesis and inhibits autophagy. Consequently, AMPK activation leads to a net balance in favour of protein degradation.

Autophagy is a conserved process from yeast to mammals for the bulk degradation and recycling of long-lived proteins and organelles^9^. Intracellular components are surrounded by double membrane-bound autophagic vesicles which then fuse with lysosomes to form autolysosomes for degradation. Controlled by autophagy-related proteins (Atgs) such as Atg5 and 7, beclin 1 and microtubule-associated protein 1 light chain 3 (LC3), autophagy plays an essential role in maintaining cellular homeostasis and was initially believed to be a non-selective process. In eukaryotic cells, autophagy occurs constitutively at low levels to facilitate homeostatic functions such as protein and organelle turnover^10^. It is rapidly upregulated when cells need to generate intracellular nutrients and energy or to remove damaging components resulting from oxidative stress or protein aggregates accumulation. Increasing evidence demonstrates that autophagy plays an essential role in cardiac remodelling to maintain cardiac function and cellular homeostasis in the heart. In certain circumstances, when excessive, the pro-survival functions of autophagy may paradoxically become deleterious^10,11^.

In this study, we dissected the signalling pathway involved in the anti-hypertrophic effect of the human β3-AR expressed in rodent cardiac myocytes, as previously used by us^3,4^. We identified β3-AR coupling to AMPK and downstream autophagy as key components for this protection.

## METHODS

### Animals

The investigation conforms to the *Guide for the Care and Use of Laboratory Animals* published by the US National Institutes of Health (NIH Publication No. 85-23, revised 1985). All experimental protocols were approved by the local Ethics Committee.

Male mice harbouring an α-MHC-driven human β3-AR transgene (β3-TG), generated as described previously^12^, were used between 12-16 weeks. Ascending aorta constriction was performed as described^4^. Briefly, after anesthetizing, a constrictive band was placed and tightened around the aorta constricted by a cannula with a width of 27G. The ligature was not tightened in sham-operated mice. Doppler measurements of trans-stenotic gradients were systematically performed at day 1, week 3 and 9 post-surgery. Only mice with a velocity higher than 2.5m/s were kept into the experiment. Mice were also submitted to the protease inhibitor leupeptin treatment to inhibit autophagic degradation (Leup, 40mg/kg, IP, 1h).

### In vitro cardiac myocytes preparations

Adult mouse ventricular myocytes (AMVM) were isolated from the hearts of 8 week-old β3-TG mice. Mice were killed by an intraperitoneal injection of sodium pentobarbital overdose (300 mg/kg) with heparin (8,000 units/kg), and the heart was rapidly excised. The ascending aorta was cannulated with a needle, and the heart was retrogradely perfused in a Langendorff perfusion system at 37°C for 5 min with perfusion buffer. This was followed by 8 min of perfusion with digestion buffer (4mg/mL trypsin, 5mg/mL liberase (Roche) and 0.3mM CaCl2). The ventricles were removed, chopped into small pieces in stop buffer (BSA 50mg/mL, 0.12mM CaCl2) and gently agitated for 3 min. The supernatant containing isolated myocytes was centrifuged (1000 rpm for 1 min), and the myocytes were resuspended in stop buffer, and subjected to a step-wise recalcification protocol (5 × 4 min stepwise increase in CaCl2 concentration from 62μM to 112μM to 212μM to 500μM to 1mM). The myocytes were plated on laminin-coated Labtek culture slides. After 1h, stop buffer was replaced by plating medium (MEM with GBS 5%, BDM 10mM, penicillin 100 U/mL and L-Glutamine 2mM).

Ventricular myocytes from 1-2 days old neonatal rats (NRVM) were isolated by collagenase/pancreatin digestion as previously described^3^. Approximately 20h post-isolation, myocytes were transferred to serum-free media and infected with a recombinant adenovirus encoding a polycistronic construct encoding the human β3-AR cDNA (form C) and GFP at a multiplicity of infection of 1.5 plaque forming units per cell; an adenovirus encoding GFP only was used as control. 24h after infection, myocytes were treated with either phenylephrine (PE, 20μM) to induce hypertrophy and/or the protease inhibitor leupeptin to inhibit autophagic degradation (Leup, 150μM) in serum-free conditions.

### AMPK knockdown by small interfering RNA transfection

NRVMs at a confluence of 50% were transfected with either control non-targeting siRNA or pooled siRNA targeting both AMPKα1 and AMPKα2 catalytic isoforms (AMPKα1/α2 siRNA) (from GE Healthcare 50nM) using lipofectamine RNAimax transfection reagent (Invitrogen) according to the manufacturer’s protocol. Phenylephrine treatment was started 66h post-siRNA transfection.

### Subcellular fractionation

Left ventricular extracts were prepared in 1 mL of 0.5M Na2CO3 (pH 11) and added to 1 mL of 80% sucrose (in 0.15M NaCl; 25mM MES, pH 6.5; final sucrose concentration, 40%). A 2-step gradient was loaded on the top of the sample (4 mL of 35% sucrose overlaid with 4 mL of 5% sucrose). After ultracentrifugation at 38000 rpm (18h, 4°C), fractions were collected from top to bottom, precipitated with trichloroacetic acid (7.2%), washed, and resuspended in Laemmli for discontinuous (8% to 12% SDS-polyacrylamide) electrophoresis and immunoblotting with antibodies (Supplemental Table 1).

### Western blotting

Denatured proteins were separated by SDS-PAGE, transferred on nitrocellulose membrane, then blocked 1h in 5% non-fat dry milk in TBS-Tween and incubated overnight in 1% milk Tween-TBS with primary antibodies. Following washing (3 × 10 min in Tween-TBS) membranes were incubated with relevant secondary antibodies in 1% milk Tween-TBS. After a final washing step (3 × 10 min Tween-TBS), membranes were revealed by chemiluminescence. Further details on primary and secondary antibodies are provided in Supplemental Table 1.

### Radiolabeled aminoacids incorporation

NRVM were plated at 1,000,000 per plate in 6-wells plates. After infection, myocytes were treated at the same time with PE (20μM) and [^14^C]l-phenylalanine (5μCi/mL) in serum-free conditions for 24h. Cells were then lysed in 50mM HEPES, 50mM KF, 1mM Kpi, 5mM EDTA, 5mM EGTA, 15mM β-mercapto-ethanol, 5% triton and incubated with 10% trichloroacetic acid for 20 min at 4°C to precipitate proteins. Precipitated material was neutralized with 100mM NaOH. The samples were precipitated a second time with 10% trichloroacetic for 20 min at 4°C. After re-suspension of protein pellets in formic acid, incorporated radioactivity was measured by liquid scintillation counting.

### Histology and immunostaining

All morphometric and histologic measurements were obtained from hearts arrested in diastole (intracardiac KCl), washed and fixed. Transversal cryo-sections were defrosted slowly in acetone, before washing (PBS; 3 × 5 min). Sections were then incubated with Wheat germ agglutinin (WGA, 1/150; 2h, in the dark). After additional washing, sections were mounted with Vectashield and observed with Axioimager Z1 Apotome system with MRm Rev3 AxioVision-Deconvolution 3D camera. Data from multiple sections per heart were analyzed using AxioVision software 4.8.2.0 (Carl Zeiss MicroImaging GmbH, Germany).

### Immunofluorescence and Proximity Ligation Assay

NRVM were fixed with 4% paraformaldehyde in PBS for 20 min. After washes (PBS, 3 × 5 min), cells were permeabilized with 0.1% triton X-100 in PBS for 10 min. After 3 washes, cells were incubated with 5% BSA in PBS for 30 min and then incubated with anti-alpha-actinin or LC3 primary antibody (Supplemental Table 1) overnight at 4°C. After 3 washes, cells were incubated with secondary antibodies diluted at 1/300 for 1h, then with Dapi 1/50000 in PBS for 5 min and mounted.

A PLA kit II (Eurogentec, Seraing, BE) was used according to the manufacturer’s instructions with slight modifications. Briefly, cardiomyocytes were fixed and transiently permeabilized (with 4% paraformaldehyde in PBS; 10 min), washed (PBS; 2 × 2 min), quenched (50mM NH4Cl; 2 × 5 min), washed, and blocked (blocking buffer; 30 min, 37°C) before being incubated overnight (4°C) with primary antibodies (Supplemental Table 1). The following day, cardiomyocytes were washed (Buffer A; 2 × 5 min) and incubated with the PLA probe secondary antibodies (plus = anti-rat (1/150 v/v); minus =anti-mouse (1/8 v/v); 60 min, 37°C). After this step cardiomyocytes were washed (Buffer A), and the ligation and amplification steps carried out by addition of ligase (1/40 v/v in ligation buffer; 30 min, 37°C) followed by washing (Buffer A), and addition of polymerase (1/80 v/v in amplification buffer; 100 min, 37°C). After a final washing step (2 × 10 min 1 × Buffer B, 1min 0.1x Buffer B), Labteks were mounted with Duolink mounting medium with DAPI, and observed with Axioimager Z1 Apotome system with MRm Rev3 AxioVision-Deconvolution 3D camera (Zeiss, Zaventem, BE). The data analysis was performed with Axiovison software (Zeiss).

### Statistical analysis

Results are expressed as mean ± SEM calculated from the measurements obtained from each group of cells and mice. When normal distribution (Kolmogorov-Smirnov test) was confirmed, raw data were analyzed using 2-way ANOVA followed by Bonferroni post-hoc test for multi-group comparisons. In absence of normal distribution, data were compared using nonparametric tests (Kruskall-Wallis followed by Dunn correction for multiple comparisons or Mann–Whitney). Statistical significance was accepted at the level of P<0.05

## RESULTS

### β3-AR and AMPK are co-localized in caveolae

We used a transgenic mouse line with cardiac myocyte-specific, moderate level expression of the human β3-AR (β3-TG) to examine the subcellular distribution of the β3-AR and AMPK. Cardiac extracts were resolved by isopycnic ultracentrifugation. As previously shown, the β3-AR protein was mainly distributed in light fractions, together with a majority of caveolin-3 (Cav-3), the main caveolar coat protein expressed in muscle cells^3^. Moreover, in the same extracts, a part of AMPK (using antibody recognizing both AMPKα1/2 catalytic subunits) co-segregated with Cav-3 signals (Figure 1A). To verify that this corresponds to the location in intact cells, we used proximity ligation assay to examine the co-localization of AMPK and Cav-3 in AMVM; as illustrated in Figure 1B, positive fluorescence signals indicated co-localisation of the 2 proteins in adult cardiomyocytes.

**Figure 1.**
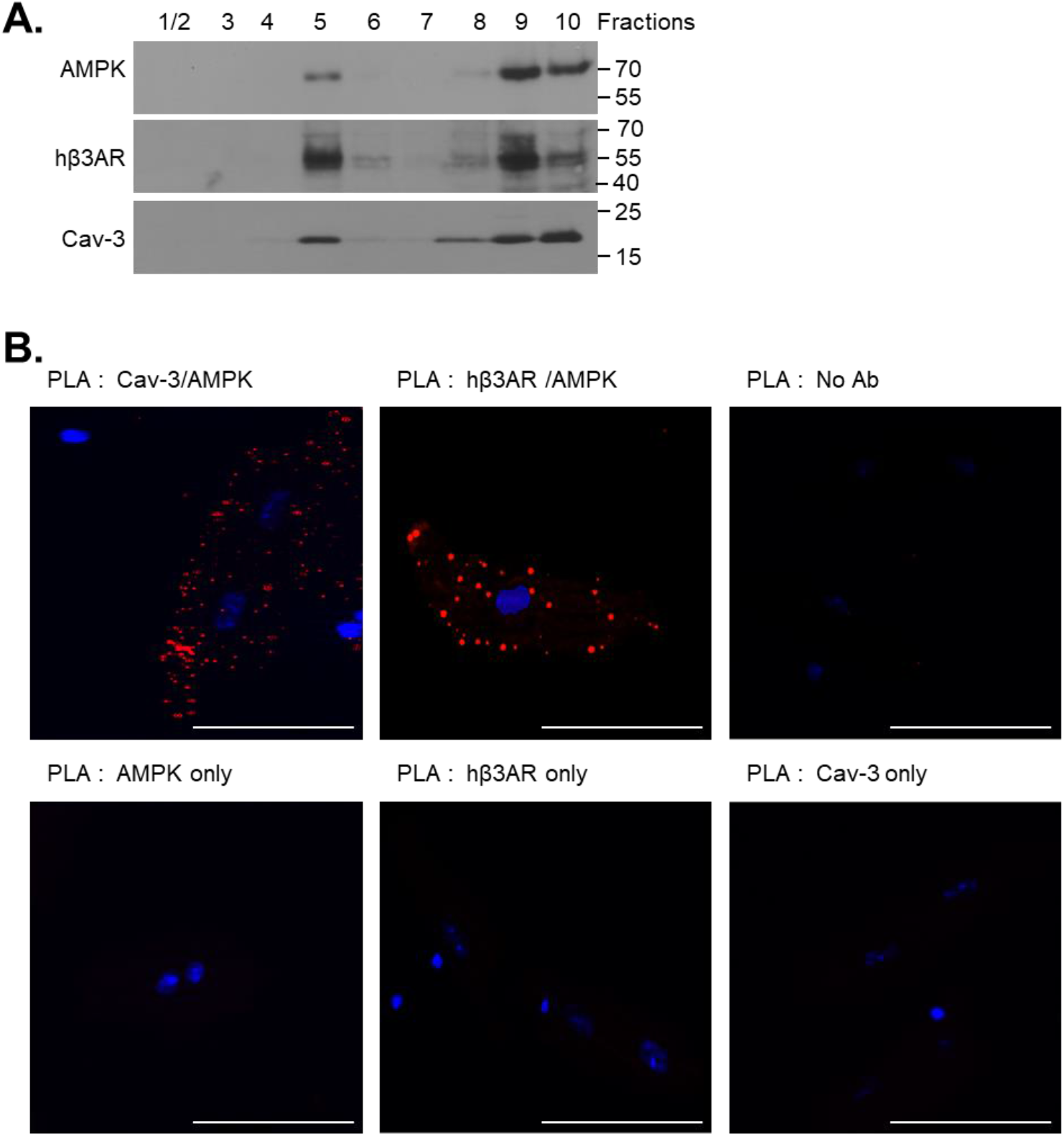
β3-adrenoceptor and AMPK subcellular localization in β3-transgenic mouse hearts. A. Subcellular fractionation of cardiac extracts of TG mice separated by ultracentifugation on sucrose gradients; each fraction (1 to 10 fractions of increasing densities) was analysed by immunoblotting for AMPK, hβ3-AR and myocyte-specific caveolin-3 (Cav-3) proteins. B. Colocalization of Cav-3 or hβ3-AR and AMPK in adult ventricular myocytes by Proximity Ligation Assay; red dots indicate co-localization of both proteins in the presence of the 2 respective primary antibodies (upper row, left and center); negative controls are illustrated in the presence of only one primary Ab or no Ab. Scale bar is 100 μm.

### Effect of neurohormonal stimuli on AMPK signalling pathway in neonatal cardiac myocytes

To examine how expression of the β3-AR could attenuate hypertrophy in response to a neurohormonal stimulus, we used an *in vitro* model of homotypic cultures of cardiac myocytes (NRVM) with adenoviral expression of the human β3-AR (AdV-β3-AR) or GFP used as control (AdV-GFP). AdV-β3-AR-infected NRVM expressed human β3-AR transcripts in proportion to the titer of recombinant viruses, as measured by RT-qPCR (Figure 2A). Immunofluorescence staining confirmed the expression of the human β3-AR protein at the membrane of infected NRVMs (Figure 2B). We also used proximity ligation assay to examine the co-localization of AMPK and β3-AR in NRVM; as illustrated in Figure 2C, positive fluorescence signals indicated co-localisation of the 2 proteins both in Adv-GFP and in AdV-β3-AR-infected neonatal cardiac myocytes, both in absence and after stimulation with phenylephrine. No PLA signal was observed in negative controls (i.e. in the absence of either primary antibody). Control, GFP-infected myocytes developed hypertrophy in response to phenylephrine (PE)^3^. This was associated with a decrease in AMPK signalling pathway, identified by a decrease in Thr172 phosphorylated-AMPK (Figure 2BD) and a decrease in phosphorylation of its substrate acetyl-CoA carboxylase (ACC) on Ser79 (Figure 2E). The inhibition of AMPK pathway correlated with the activation of mTOR pathway, assessed from the phosphorylation state of its indirect downstream target, the ribosomal S6 protein (S6) on Ser235/236 (Figure F). This was associated with an increase in radiolabelled amino acids integration into proteins (Figure 2G). In contrast, AdV-β3-AR infected myocytes retained basal level of AMPK, ACC and S6 phosphorylation and unchanged protein synthesis rate despite PE stimulation (Figure 2D-G).

**Figure 2.**
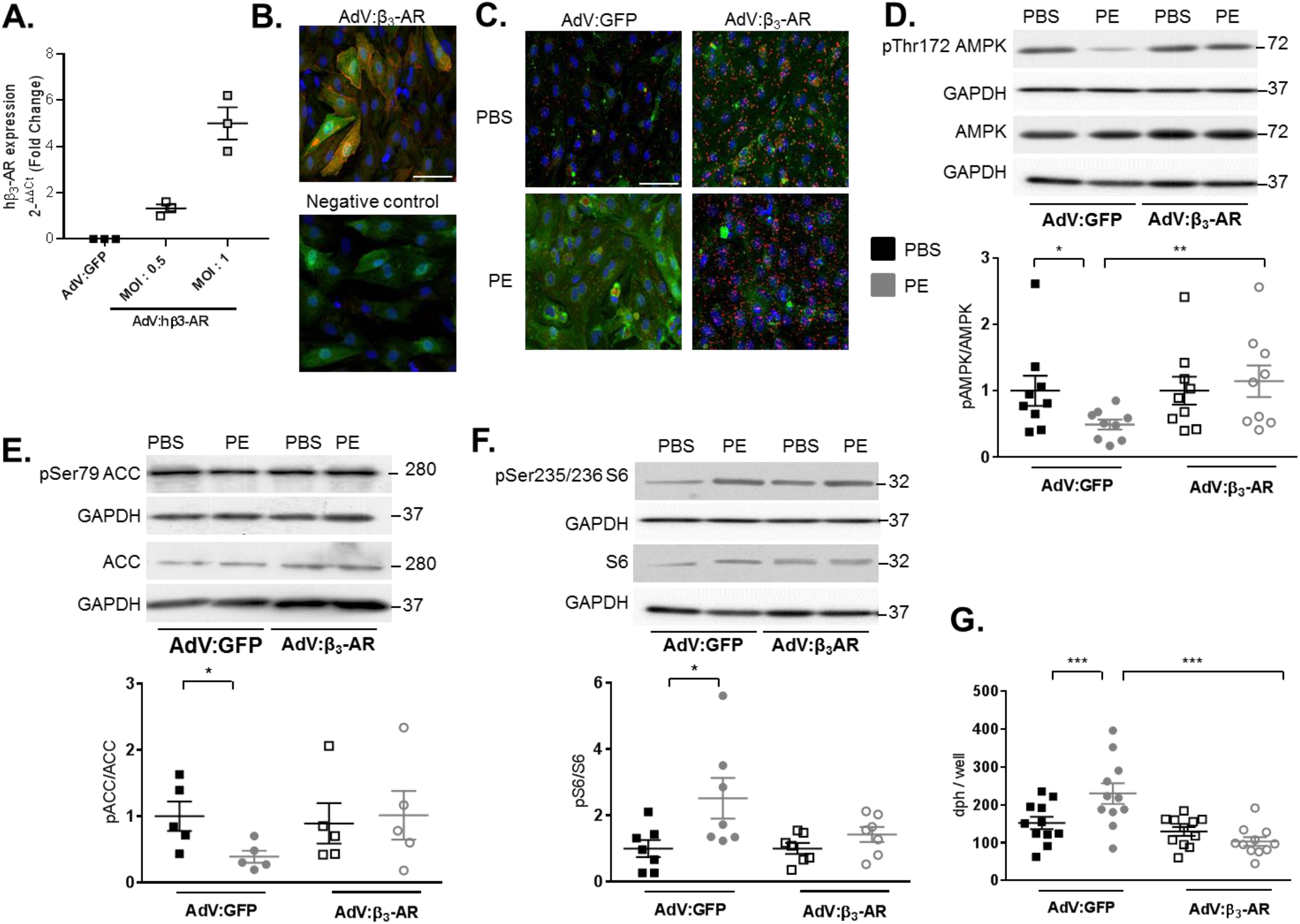
Effect of β3-adrenoceptor expression on AMPK signaling pathway after phenylephrine treatment in neonatal cardiac myocytes. A: RT-qPCR of hβ3-AR mRNA in AdV-GFP (control) and AdV-β3-AR-infected neonatal rat ventricular myocytes (NRVM). B: (Upper) Immunocytochemical detection of hβ3-AR in AdV-β3-AR-infected NRVM; (Lower) negative control in absence of primary antibody. C: Colocalization of hβ3-AR and AMPK in AdV-GFP (control) and AdV-β3-AR-infected NRVM by Proximity Ligation Assay; red dots indicate co-localization of both proteins in the presence of the 2 respective primary antibodies. Scale bar is 50 μm. (Negative controls in absence of either primary antibody are in Fig S1). D, E, F: Western blot of phosphorylated AMPK (D), ACC (E) and S6 (F) in response to phenylephrine (PE) in AdV-GFP (control) and AdV-β3-AR-infected NRVM (immunoblotted signals normalized to respective control values in PBS) G: Quantification of radiolabelled phenylalanine incorporated into proteins in response to PE in AdV-GFP (control) and AdV-β3-AR-infected NRVMs.*, P<0.05; ** P<0.01; ***P<0.001 from at least 5 experiments.

### Effect of haemodynamic overload on AMPK signalling pathway in mice

Adult β3-TG mice were subjected to transaortic constriction (TAC) and their phenotype analysed at 9 weeks post-TAC. Morphometric data are illustrated in Figure 3A. All parameters were identical between genotypes in sham-operated mice. All mice included in the TAC study (β3-TG and WT) developed a trans-stenotic gradient with maximum velocity (by Doppler echo) of at least 2.5 m/s, and gradients were comparable between genotypes. WT mice developed myocardial hypertrophy, as evidenced by an increase in left ventricular weight (LVW) normalized to tibial length (TL) (Figure 3A) and increased myocyte transverse area measured histologically (Figure 3B). However, increases in both parameters were significantly milder in β3-TG mice.

**Figure 3.**
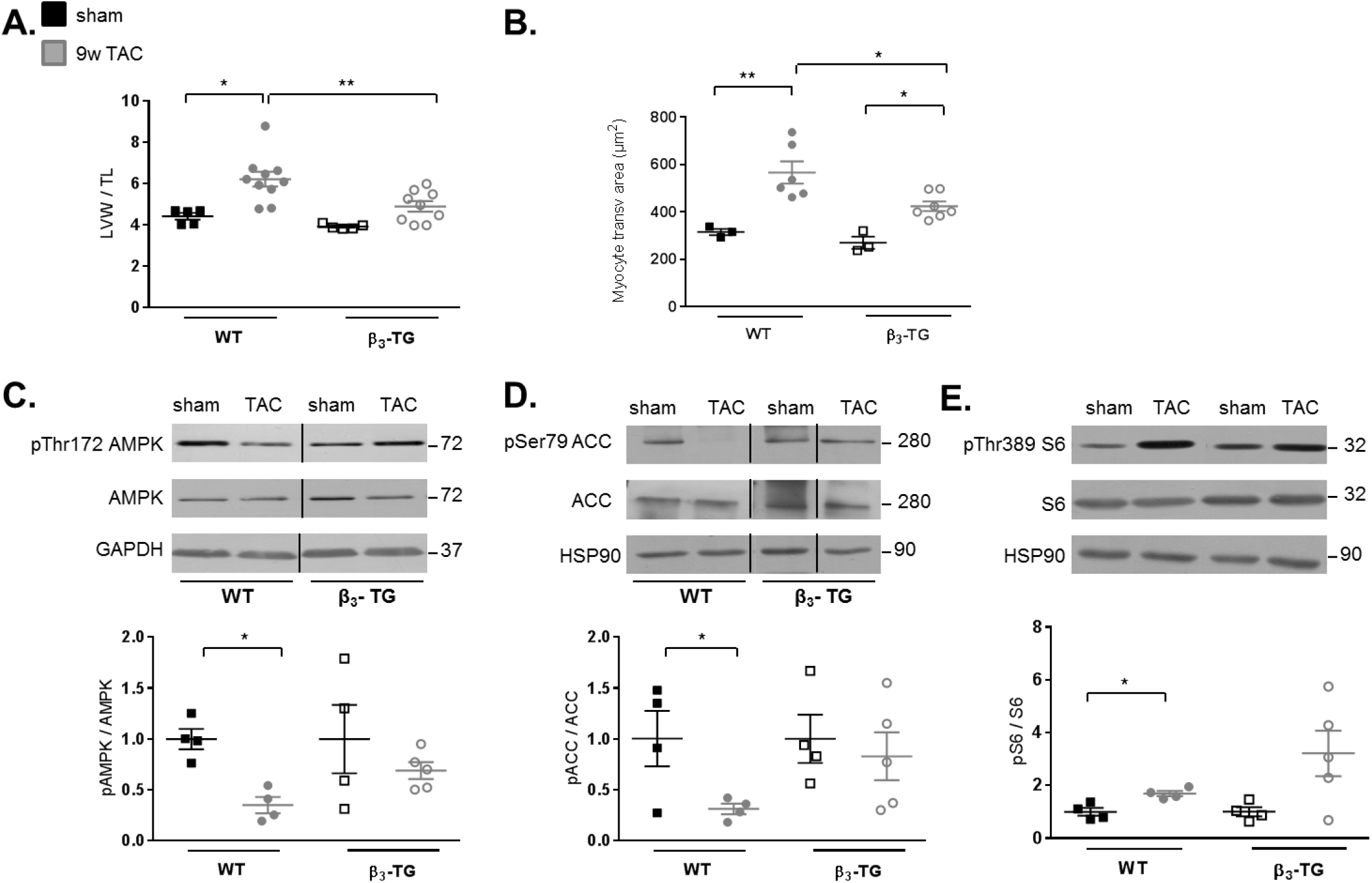
Effect of β3-adrenoceptor expression on AMPK signaling pathway after 9 weeks of TAC in mice. A,B: Changes in (A) LV mass assessed by gravidometry, (B) myocyte transverse section areas at 9 weeks post TAC in WT and β3-TG mice. C,D, E: Western blot of phosphorylated AMPK (C), ACC (D) and S6 (E) at 9 weeks post TAC in WT and β3-TG mice (signal normalized to respective control values in sham).*, P<0.05; ** P<0.01; from at least 3 sham mice and 6 TAC mice.

At the molecular level, 9 weeks of TAC resulted in a significant decrease in AMPK signalling, as identified by a decrease in AMPK phosphorylation on Thr172 (Figure 3C), a decrease in ACC phosphorylation on Ser79 (Figure 3D) and an increase in S6 phosphorylation on Thr389 (Figure 3E) in WT mice. However, basal levels of phospho-Thr172-AMPK (Figure 3C), phospho-Ser79-ACC (Figure 3D) and phospho-Thr389-S6 (Figure 3E) appeared unchanged, albeit with some variability, from sham-operated controls in β3-TG mice.

### Role of AMPK in the anti-hypertrophic effect of β3-AR

These data and our observation of co-localization of β3-AR and (at least part of) AMPK in caveolar fractions (see above), prompted us to examine the causality of AMPK in the modulation of the hypertrophic response by β3-AR. To do this, we tested the effect of AMPK downregulation on the hypertrophic response to PE in NRVM. Cultured NRVMs with adenoviral expression of the human β3-AR (or GFP control) were transfected with siRNAs targeting both AMPKα1/2 catalytic subunits (or scramble control) and then treated with PE. High efficiency of AMPKα1/2 knockdown was validated by immunoblot showing the complete disappearance of AMPKα signal (Figure 4). As expected, β3-AR expression almost completely attenuated the hypertrophic response to PE compared to GFP control in si-scramble-transfected cells (Figure 4, bottom left). AMPK silencing potentiated the hypertrophic response to PE in GFP-expressing myocytes; although β3-AR expression still attenuated the hypertrophic response to PE, it was only reduced by half, in si-AMPK-transfected cells compared to GFP control (Figure 4, bottom right).

**Figure 4.**
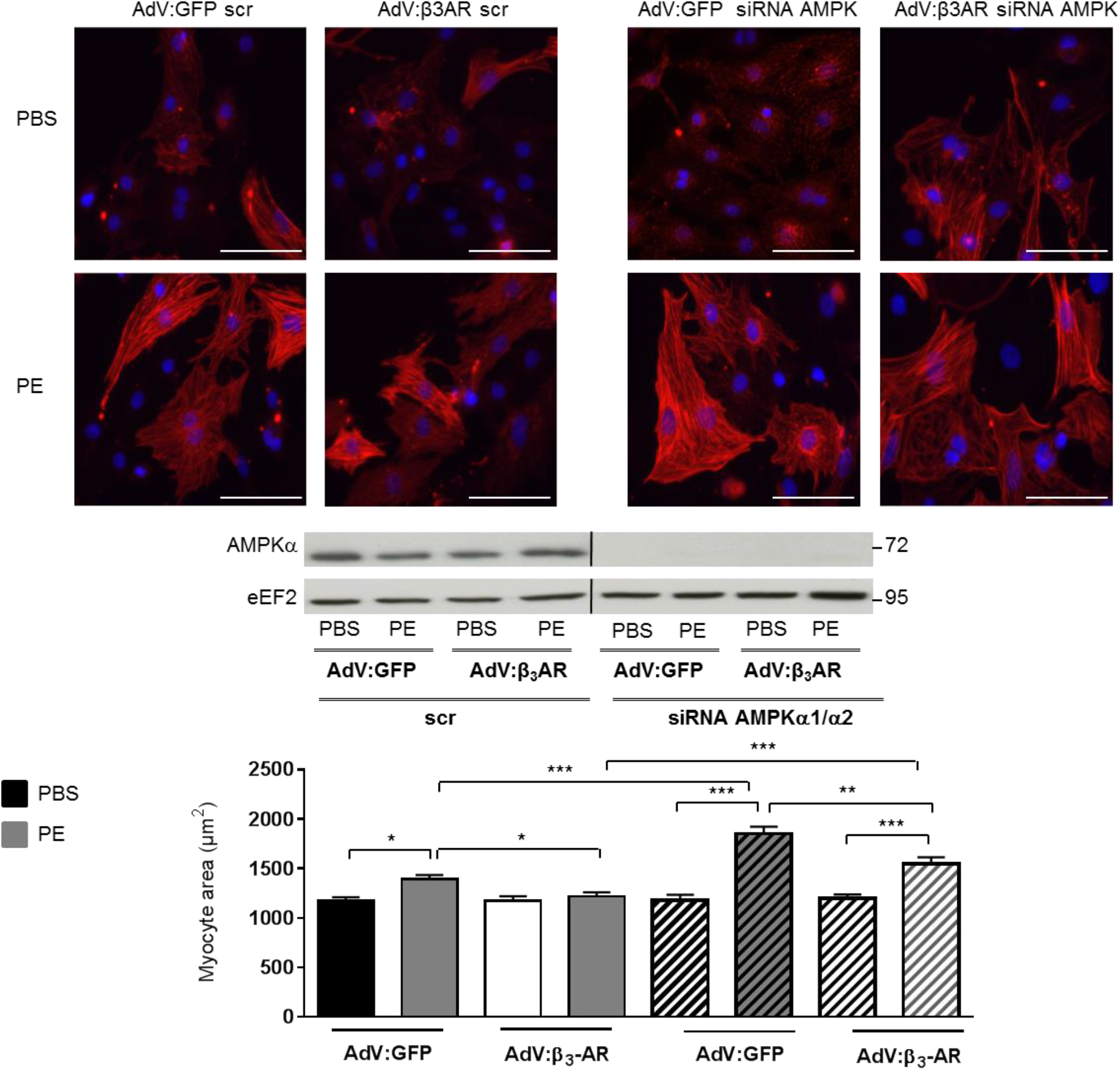
Effect of AMPK inhibition on β3-adrenoceptor regulation of hypertrophy in cardiac myocytes. Changes in cell size in AdV-GFP and AdV-β3-AR infected NRVMs after PE alone or with concurrent administration of siRNA targeting AMPK-α1/2 (or scramble control). Upper: representative images of NRVM stained for alpha-actinin (red) and nuclei (DAPI: blue). Scale bar is 100 μm. Lower: Representative Western blot showing the inhibitory effect of the siRNA construct on AMPK α expression. Bottom: Bar graphs illustrating the quantification of myocyte sizes under the different treatment conditions *, P<0.05; ** P<0.01; ***, P<0.001; from 6 experiments (at least 185 cells analyzed).

### Effect of neurohormonal stimuli on autophagy signalling pathway in cardiac myocytes

Control AdV-GFP infected myocytes developed hypertrophy in response to PE together with a decrease in autophagy identified by a decrease in LC3 II/I ratio (Figure 5A), an increase in p62 expression (Figure 5B) and a 24.5% decrease of autophagic cells (by LC3 immunostaining quantification) (Figure 5C). By contrast, AdV-β3-AR infected myocytes maintained their basal level of autophagy (monitored with the same 3 criteria) despite PE stimulation. Of note, a similar decrease in LC3 II/I ratio was observed after TAC in myocardial extracts from WT mice; however, basal levels were again maintained in β3-TG mice after TAC (Figure 5D).

**Figure 5.**
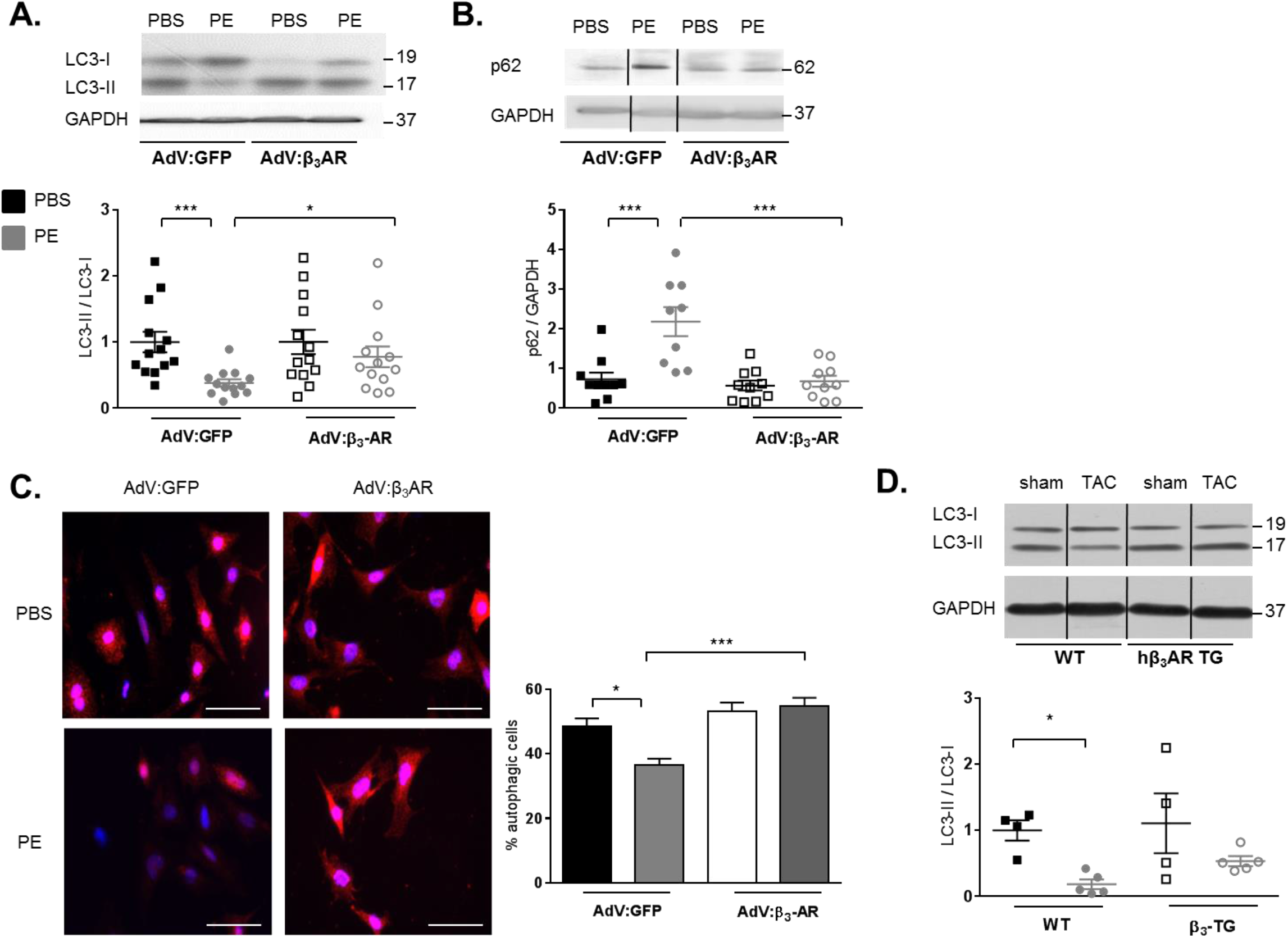
Effect of β3-adrenoceptor expression on autophagy after PE treatment in cardiac myocytes. A,B: Western blot of LC3-II/LC3-I (A) and p62 (B) in AdV-GFP and AdV-β3-AR-infected NRVMs with/without PE stimulation (signal normalized to respective control values in PBS); C: Quantification of percentage of autophagic cells in AdV-GFP and AdV-β3-AR-infected NRVMs with/without PE stimulation (at least 70 images analysed/condition); D: Western blot of LC3-II/LC3-I in cardiac extracts from WT and β3-TG mice after 9 weeks of TAC or sham operation (signal normalized to respective control values in sham).*, P<0.05; ** P<0.01; ***, P<0.001; from at least 9 experiments *in vitro* and 4 sham mice and 5 TAC mice.

We then analysed autophagic flux by inhibiting autophagic degradation with the protease inhibitor leupeptin (Figure 6). As expected, leupeptin treatment increased the level of LC3 II/I ratio and percentage of LC3-marked cells in AdV-GFP infected myocytes, unravelling a basal autophagic flux (Figure 6A and C). Leupeptin also reversed the decrease of these indexes induced by PE treatment (Figure 6A and C). However, leupeptin had no effect in AdV-hβ3 infected myocytes, suggesting that autophagy is already maximal in these cells, with or without PE (Figure 6B). Interestingly, these effects of leupeptin were reproduced in hearts of WT and β3-TG mice (Figure 6D) after i.p. injection. Similar effects were reproduced with chloroquine that inhibits the autophagososme-lysosome fusion and degradation (not shown).

**Figure 6.**
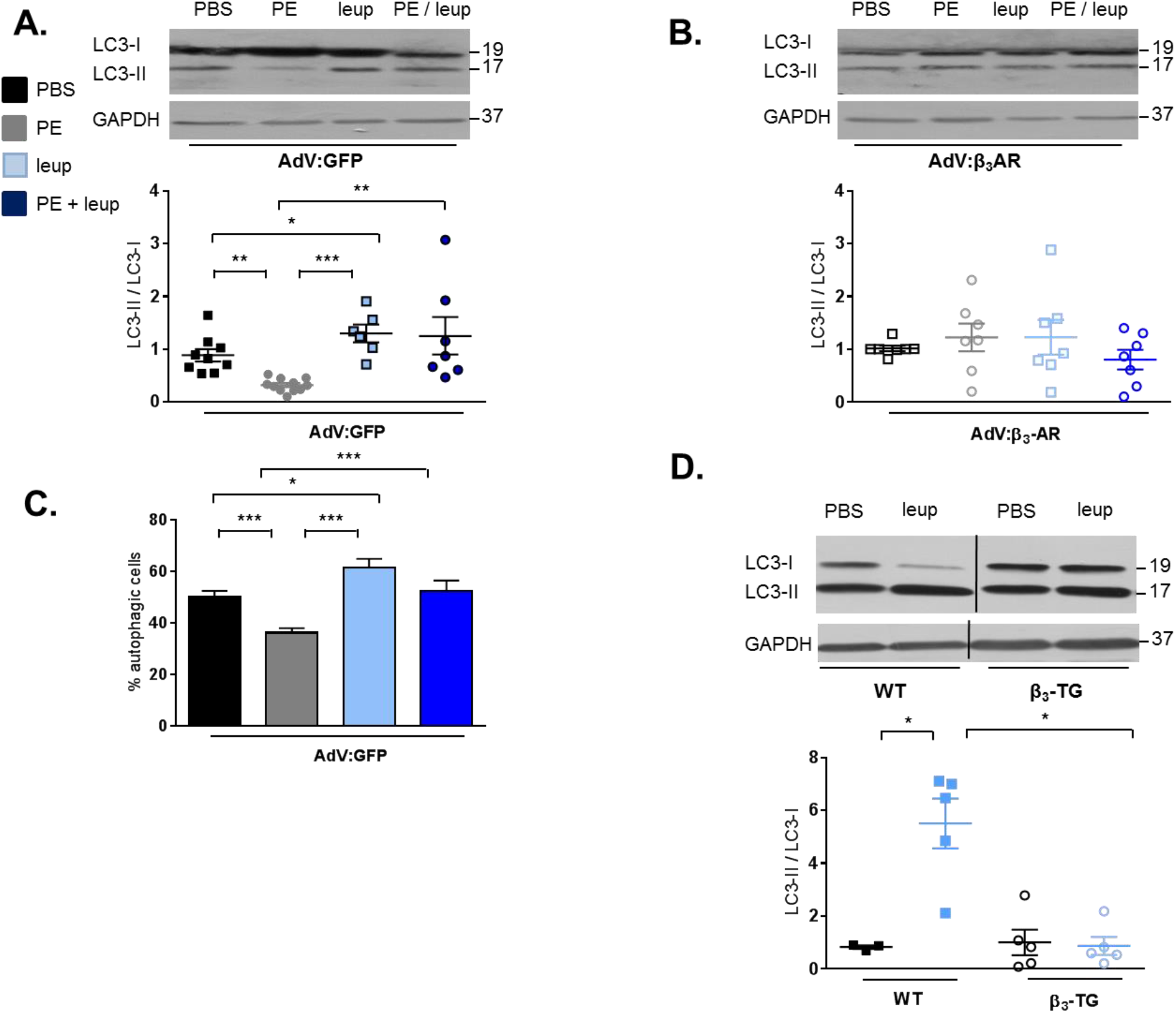
Autophagic flux in β3-adrenoceptor-expressing cardiac myocytes *in vitro* and *in vivo*. A,B: Western blot of LC3-II and LC3-I in response to PE with/without leupeptin (leup) in AdV-GFP and AdV-β3-AR-infected NRVMs (signal normalized to respective values in PBS control); C: Quantification of percentage of autophagic cells in response to PE with/without leupeptin in AdV-GFP infected NRVMs (at least 33 images analysed/condition); D:Western blot of LC3-II and LC3-I in response to leupeptin in cardiac extracts from WT and β3-TG mice (signal normalized to respective values in PBS control).*, P<0.05; ** P<0.01; ***, P<0.001; from at least 6 experiments in vitro and 4 mice in each group.

**Figure 7.**
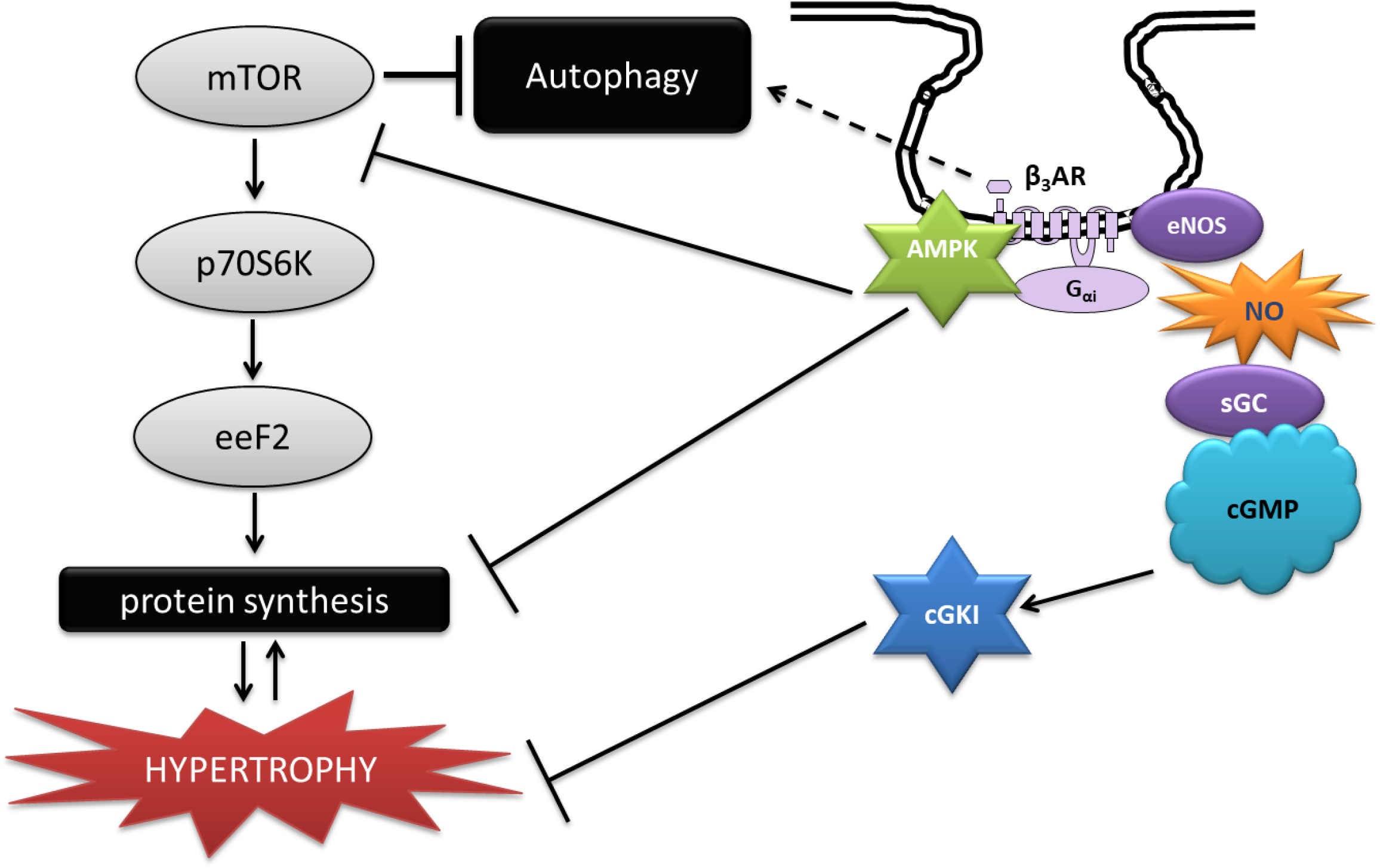
Schematic representation of β3-adrenoceptor inhibitory signalling to hypertrophic pathways. β3-AR colocalizes with several effectors in caveolar membranes of cardiac myocytes; (Right) previously identified coupling to nitric oxide synthase (NOS) activates NO/sGC/cGMP/cGK-I pathway known to attenuate stress-induced hypertrophy (for review, see Farah et al. Nat Rev Cardiol 2018;15(5):292-316); (Left) the presently identified interaction with AMPK recruits this pathway to inhibit protein synthesis and hypertrophy under stress as well as the upstream pro-hypertrophic mTOR complex; inhibition of mTOR would also relieve its negative effect on autophagy, thereby increasing autophagic flux, although the latter may also be independently activated by β3-AR coupling to alternative effectors.

## DISCUSSION

The main findings of this study are as following; i. the β3-AR co-localizes with (part of) AMPK in caveolae-enriched membrane fractions of AMVM; ii. β3-AR expression restores basal level of AMPK signaling and autophagy in response to neurohormonal (PE) and haemodynamic (TAC) stress; iii. AMPK is, at least partially, involved in the anti-hypertrophic effect of β3-AR.

The β3-AR was first cloned from a human adipocyte cDNA library and is expressed mainly in brown/beige adipocytes, where it mediates non–shivering thermogenesis^13^. It was later found to be expressed in cardiac myocytes from several species, including human ventricular cardiac biopsies in which, contrary to β1-2ARs, β3-AR stimulation decreases contractile tone *in vitro*^14^. As β3-AR expression increases in cardiomyopathic hearts ^15^, this led to the hypothesis that β3-AR may confer protection from the deleterious overstimulation of β1-2AR as observed in the course of heart failure progression^16,17^. In this study, we used a heterozygous transgenic mouse with cardiac-specific, moderate expression of the human β3-AR that replicated the expression level of the receptor and the functional response to β3-AR agonists observed in human biopsies^4,12^. Contrary to β1-2 AR transgenics, these β3-AR transgenic mice are protected from neurohormone and TAC-induced hypertrophic remodelling^3^. The present study analysed the precise mechanisms underlying this protection.

We previously reported that the recombinant β3-AR co-localized with caveolin-3 in lipid rafts/caveolae^3^, together with the endothelial nitric oxide synthase (eNOS)^17^. Here, we show that AMPK is also partially expressed in caveolae as previously observed in rat cardiac myocytes^18^.

Since adult cardiomyocytes have little capacity for cellular proliferation, their only means of growth is hypertrophy. Cardiac hypertrophy is considered to be an initially adaptive response, counteracting increased wall tension and helping to maintain cardiac output. Cardiac mass is determined by the balance between protein synthesis and degradation, known to be tightly regulated during the development of cardiac hypertrophy^19^. Accordingly, we observed that hypertrophy induced a decrease in phosphorylation of AMPK on its Thr172 activation site, resulting in AMPK signaling pathway inhibition. Modulation of AMPK activity during cardiac hypertrophy is controversial in the literature. Indeed, in one study, no modification in AMPK activity or phosphorylation was found during the first 8 weeks of TAC-induced hypertrophy^20^ whereas another study reported an activation after 17 to 19 weeks post-surgery^21^. Such late AMPK activation can be hypothesized as a negative feedback mechanism to reduce hypertrophy progression. In both *in vitro* and *in vivo* protocol settings used in our study, hypertrophic stimuli clearly induced AMPK inhibition. Hypertrophy-mediated AMPK downregulation was associated with a decrease in S6 phosphorylation and an increase in radiolabelled aminoacid incorporation into proteins. Expression of the human β3-AR clearly restored AMPK signaling pathway, and attenuated hypertrophy in PE-stimulated NRVMs and hearts with TAC-induced hypertrophic remodelling. The implication of AMPK downstream β3-AR activation was clearly demonstrated *in vitro* with the use of specific siRNA against AMPK catalytic subunits, which partly restored the hypertrophic response in β3-AR overexpressing cells. This is in line with recent data showing that exercise training protects the heart against ischemia/reperfusion injury by activating eNOS via the stimulation of a β3-AR-AMPK signaling pathway^22^. Note that we previously demonstrated that the heterologously expressed β3-AR is constitutively active^3^, explaining its tonic effect in absence of stimulation with specific agonists. Nevertheless, despite AMPK silencing, the β3-AR retained a diminished, but still present anti-hypertrophic effect, that could be explained by alternative coupling to pathways downstream endothelial (e-) and neuronal (n-) nitric oxide synthases (NOS), as previously reported by us^3^.

mTOR integrates upstream signals from various stresses such as cellular energy levels and inhibits autophagy. Reciprocally, AMPK could activate autophagy by inhibiting mTOR and both AMPK and mTOR can themselves be phosphorylated by Unc-51-like autophagy activating kinase 1 (ULK1), providing an additional level of regulatory feedback to modulate autophagy^23^. Autophagy is an evolutionarily conserved mechanism that targets damaged proteins or organelles for degradation through lysosomes. Autophagy can be non selective or organelle-specific, such as mitochondria-specific autophagy and plays an essential role in maintaining cellular homeostasis in the heart at baseline and in response to stress. Indeed, reduced autophagy gradually accumulates harmful proteins and generates reactive oxygen species from damaged mitochondria in the hypertrophied hearts. In line with this, we observed that TAC- or PE-induced hypertrophy was accompanied with a decrease in autophagy both *in vitro* and *in vivo* and expression of the human β3-AR restores autophagy, consistent with its protective effect against hypertrophy and remodelling. We used leupeptin, an inhibitor of autophagolysosomal proteases, (as well as chloroquine, an inhibitor autophagosome-lysosome fusion and degradation) to monitor autophagic flux in these models. Leupeptin has already been reported to induce LC3II accumulation in adult rat cardiomyocytes^24^. As expected, we observed an accumulation of LC3II in NRVMs and hearts treated with these inhibitors. Interestingly, leupeptin (or chloroquine) did not further increase LC3II in β3-AR-expressing myocytes *in vitro* or *in vivo*, possibly because autophagy flux may already have been maximally stimulated.

These results are consistent with previous observations on autophagy in cardiac hypertrophy. Indeed, autophagy was shown to be suppressed in TAC-induced hypertrophied hearts at 1 week post-surgery compared to sham-operated hearts^25^ or diminished in response to supravalvular aortic constriction^26^ or β-adrenergic stimulation^27^. Interestingly, autophagy has been shown to participate in the anti-hypertrophic action of AMPK^28^. Nevertheless, we cannot exclude that β3-AR may activate autophagy independently of AMPK at this point. More generally, autophagy is also decreased in the aging heart whereas cardiovascular diseases progressively increase with age^29^. Increasing evidence suggests that autophagy is involved in the regulation of lifespan and aging. Autophagy may be especially important in non-proliferating cells, such as differentiated cardiac myocytes, in which there is no dilution of accumulated toxic material with cell division.

### Limitations and perspectives

Admittedly, our observations are based on models with heterologous expression of a recombinant human β3-AR in cardiac myocytes *in vitro* and *in vivo*, with the caveat that genetically reconstituted signalling may not faithfully replicate the coupling of the endogenous receptor *in situ*. However, we used a line of transgenic animals selected on the basis of levels of β3-AR expression that replicates the abundance observed in human myocardial biopsies, as demonstrated previously^4^. Given the ubiquitous expression and widespread role of AMPK, it would be of interest to verify similar coupling to β3-AR in other tissues. β3-AR transcripts^30^ and proteins^31^ have clearly been identified in human white and brown adipocytes, where they can be activated by selective β3-AR agonists, such as mirabegron^30^. Given the emerging importance of brown adipose tissue in regulating metabolism and cardiovascular function^32,33^, and the role of AMPK in promoting mitochondrial biogenesis^34^, critical for brown fat lipolytic activity, we can speculate that the β3-AR coupling mechanism identified here could be exploited therapeutically to prevent or treat cardiovascular diseases associated with obesity and disordered metabolism. Such systemic effects are currently being studied in the European BETA3-LVH trial testing the re-purposing of mirabegron in patients with structural cardiac remodelling with or at risk of developing heart failure with preserved ejection fraction^35^. Further, β3-AR modulation of visceral adipose tissue may have remote effects on cardiac aging by regulating fibroblast senescence^36^, although this has not been directly demonstrated so far.

In conclusion, moderate expression of β3-AR in cardiac myocytes protects from myocardial remodelling by modulation of AMPK signaling pathway and autophagy. These observations may open new therapeutic perspectives with specific β3-AR agonists for the treatment of hypertrophic cardiomyopathies.

## ACKNOWLEDGEMENTS

We thank Delphine De Mulder and Lucie Caërou for excellent technical work.

## FUNDINGS

Work funded by grants from the Fonds National de la Recherche Scientifique (FNRS; PDR T.0144.13), the Federation Wallonie-Bruxelles (Action de Recherche Concertée ARC11-16/035) and European Union (UE LSHM-CT-05-018833) to JLB. EDD was a Marie-Curie Fellow of the European Commission. RG received funding support from Fonds de Recherche Clinique, Université catholique de Louvain, Belgium. CB is Postdoctorate Clinical Master Specialist of the FNRS, Belgium. LB is Senior Research Associate of the FNRS, Belgium.

## CONFLICT OF INTEREST

none declared

---

**Figure S1.**
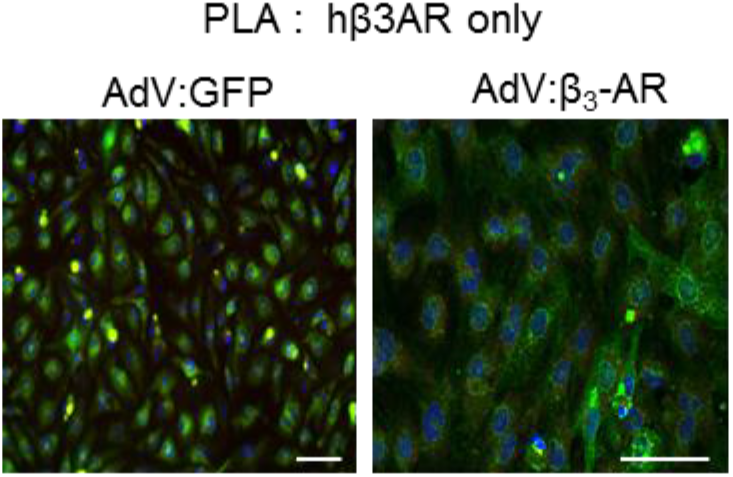
Negative control of colocalization of hβ3-AR and AMPK. Negative control in absence of primary antibody for AMPK for colocalization of hβ3-AR and AMPK in AdV-GFP (control) and AdV-β3-AR-infected NRVM by Proximity Ligation Assay. Scale bar is 50 μm.

**Supplemental Table 1:**
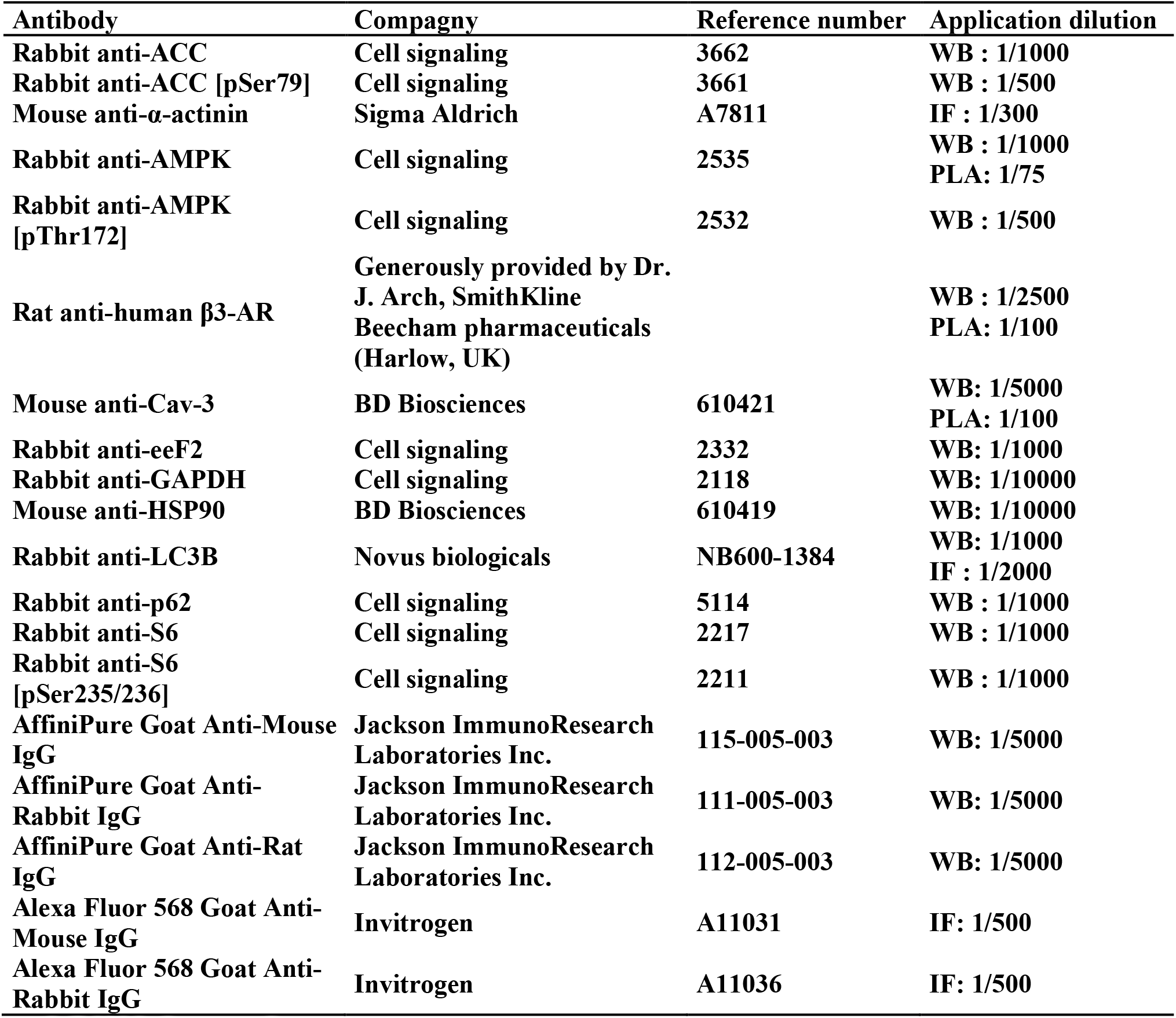
List of antibodies used.

| **Antibody** | **Compagny** | **Reference number** | **Application dilution** |
| --- | --- | --- | --- |
| **Rabbit anti-ACC** | **Cell signaling** | **3662** | **WB : 1/1000** |
| **Rabbit anti-ACC [pSer79]** | **Cell signaling** | **3661** | **WB : 1/500** |
| **Mouse anti-α-actinin** | **Sigma Aldrich** | **A7811** | **IF : 1/300** |
| **Rabbit anti-AMPK** | **Cell signaling** | **2535** | **WB : 1/1000**<br>**PLA: 1/75** |
| **Rabbit anti-AMPK [pThr172]** | **Cell signaling** | **2532** | **WB : 1/500** |
| **Rat anti-human β3-AR** | **Generously provided by Dr. J. Arch, SmithKline Beecham pharmaceuticals (Harlow, UK)** |  | **WB : 1/2500**<br>**PLA: 1/100** |
| **Mouse anti-Cav-3** | **BD Biosciences** | **610421** | **WB: 1/5000**<br>**PLA: 1/100** |
| **Rabbit anti-eeF2** | **Cell signaling** | **2332** | **WB: 1/1000** |
| **Rabbit anti-GAPDH** | **Cell signaling** | **2118** | **WB: 1/10000** |
| **Mouse anti-HSP90** | **BD Biosciences** | **610419** | **WB: 1/10000** |
| **Rabbit anti-LC3B** | **Novus biologicals** | **NB600-1384** | **WB: 1/1000**<br>**IF : 1/2000** |
| **Rabbit anti-p62** | **Cell signaling** | **5114** | **WB : 1/1000** |
| **Rabbit anti-S6** | **Cell signaling** | **2217** | **WB : 1/1000** |
| **Rabbit anti-S6 [pSer235/236]** | **Cell signaling** | **2211** | **WB : 1/1000** |
| **AffiniPure Goat Anti-Mouse IgG** | **Jackson ImmunoResearch Laboratories Inc.** | **115-005-003** | **WB: 1/5000** |
| **AffiniPure Goat Anti-Rabbit IgG** | **Jackson ImmunoResearch Laboratories Inc.** | **111-005-003** | **WB: 1/5000** |
| **AffiniPure Goat Anti-Rat IgG** | **Jackson ImmunoResearch Laboratories Inc.** | **112-005-003** | **WB: 1/5000** |
| **Alexa Fluor 568 Goat Anti-Mouse IgG** | **Invitrogen** | **A11031** | **IF: 1/500** |
| **Alexa Fluor 568 Goat Anti-Rabbit IgG** | **Invitrogen** | **A11036** | **IF: 1/500** |

